# Mapping human laryngeal motor cortex during vocalization

**DOI:** 10.1101/2020.02.20.958314

**Authors:** Nicole Eichert, Daniel Papp, Rogier B. Mars, Kate E. Watkins

**Affiliations:** Wellcome Centre for Integrative Neuroimaging, Centre for Functional MRI of the Brain (FMRIB), Nuffield Department of Clinical Neurosciences, John Radcliffe Hospital, University of Oxford, Oxford, United Kingdom; Donders Institute for Brain, Cognition and Behaviour, Radboud University Nijmegen, Nijmegen, The Netherlands; Wellcome Centre for Integrative Neuroimaging, Department of Experimental Psychology, University of Oxford, Oxford, United Kingdom

**Keywords:** Functional MRI, Larynx, Myelin, Quantitative MRI, Speech

## Abstract

The representations of the articulators involved in human speech production are organized somatotopically in primary motor cortex. The neural representation of the larynx, however, remains debated. Both a dorsal and a ventral larynx representation have been previously described. It is unknown, however, whether both representations are located in primary motor cortex. Here, we mapped the motor representations of the human larynx using fMRI and characterized the cortical microstructure underlying the activated regions. We isolated brain activity related to laryngeal activity during vocalization while controlling for breathing. We also mapped the articulators (the lips and tongue) and the hand area. We found two separate activations during vocalization – a dorsal and a ventral larynx representation. Structural and quantitative neuroimaging revealed that myelin content and cortical thickness underlying the dorsal, but not the ventral larynx representation, are similar to those of other primary motor representations. This finding confirms that the dorsal larynx representation is located in primary motor cortex and that the ventral one is not. We further speculate that the location of the ventral larynx representation is in premotor cortex, as seen in other primates. It remains unclear, however, whether and how these two representations differentially contribute to laryngeal motor control.

## 1. Introduction

The voluntary control of highly complex speech movements is regarded as one aspect of behavior that is unique to humans (Fitch 2017). Speech production requires the fine coordination of a large number of muscles to control the supralaryngeal articulators, respiration, and the vocal folds in the larynx during voice production (Jürgens 2002). In addition to its role as main source of vocal sound production, the larynx is implicated in various biological functions such as protection of the airways and swallowing (Ludlow 2005). Several pairs of intrinsic and extrinsic muscles connect the laryngeal cartilages with each other and to the skeleton. During vocalization, these muscles are controlled in a complex fashion, so that the tension in the vocal folds allows them to be set into vibration as air from the lungs passes through them. Several lines of evidence showed that the ventral part of the precentral gyrus and the central sulcus in the human brain are involved in speech motor control (Bohland and Guenther 2006; Ackermann et al. 2014). The question as to which brain areas specifically control laryngeal activity during vocalization, however, remains debated.

Nearly 100 years ago, direct cortical stimulation of the ventral portion of the precentral gyrus in the human brain was shown to elicit vocalization (Foerster 1936; Penfield and Boldrey 1937). In more recent times, functional brain imaging studies show highly inconsistent results when mapping laryngeal activity during vocalization (reviewed in Belyk & Brown, 2017). Several studies report activity evoked by vocalization in both a ventral portion of the precentral gyrus located close to the Sylvian fissure and in a more dorsal portion of central sulcus and precentral gyrus (Terumitsu et al. 2006; Galgano and Froud 2008; Olthoff et al. 2008; Grabski et al. 2012). Some studies report vocalization-evoked activity in the dorsal location only (Sörös et al. 2006; Brown et al. 2008; Kleber et al. 2013; Belyk and Brown 2014; Belyk et al. 2018). Only a few studies specifically isolated laryngeal activity during voice production by contrasting it with supralaryngeal articulation (Sörös et al. 2006; Terumitsu et al. 2006; Brown et al. 2009; Grabski et al. 2012), but the results were inconsistent across studies.

Another line of evidence comes from direct neurophysiological recordings from the cortical surface using implanted high-density electrode arrays in patients being prepared for epilepsy surgery. These studies also show activity in both a dorsal and a ventral region during speech production (Bouchard et al. 2013; Chang et al. 2013; Toyoda et al. 2014; Breshears et al. 2015; Dichter et al. 2018). Interpretation of electrocorticography (ECoG) studies, however, is inherently limited due to the typically small sample size and potential neurological abnormalities in the patients’ brains.

In addition to these inconsistent results, an additional confound in several studies mentioned above is breathing. Voice production and respiration functionally interact during speech production and volitional expiration has been shown to activate motor areas that are located close to the putative dorsal laryngeal representation (Ramsay et al. 1993; Evans et al. 1999; McKay et al. 2003). Several previous studies, however, did not control for breathing (e.g. Brown et al., 2008; Sörös et al., 2006). Those studies that specifically studied vocalization and breathing found largely overlapping activity during both conditions and a contrast showed no difference in motor cortex indicating that the activity was not specific to larynx activity (Loucks et al. 2007; Simonyan et al. 2009).

In addition to a lack of control for confounds such as breathing and supralaryngeal articulation, most neuroimaging studies mentioned above did not assess individual differences in the activation patterns. It is common to report group-level cluster-corrected results following volumetric nonlinear image registration of task-activation maps. Interpreting group-level results, however, might obscure subject-specific features and inter-individual variability, which limits sensitivity and functional resolution (Bennett and Miller 2010; Nieto-Castañón and Fedorenko 2012; Bouchard et al. 2013; Woo et al. 2014). This lack of individual detail might have caused a failure to detect one of the cortical larynx representations in previous studies. Moreover, averaging of small sample sizes with high variability, as often performed in ECoG studies, can show two distinct larynx representations, even when individual patients show activity in only one of the regions (Bouchard et al. 2013).

Comparisons of human brains and those of other primates indicate strong species differences in the neural organization underlying laryngeal control during vocalization (Ackermann et al. 2014; Simonyan 2014; Kumar et al. 2016). Most notably, the location of the proposed human dorsal larynx representation in primary motor cortex is more dorsal-posterior than the non-human primate homolog, which is in a ventral premotor cortex area (Leyton and Sherrington 1917; Hast et al. 1974; Jürgens 1974; Simonyan and Jürgens 2002; Coudé et al. 2011).

These various findings have led to the proposal of an evolutionary ‘duplication and migration’ hypothesis that the larynx motor cortex comprises two structures located dorsally and ventrally in the human brain (Belyk and Brown, 2017; Jarvis, 2019; reviewed in Mars et al., 2018). This theory proposes that the dorsal larynx representation is unique to humans and that it evolved in primary motor cortex due to our especially high demands on laryngeal motor control. The ancestral primate larynx representation in premotor cortex is presumed to be homologous to the human ventral larynx representation, which migrated posteriorly, potentially into human primary motor cortex.

The cortical areas underlying the larynx representations, however, are currently unknown and it has not been tested if the ventral larynx representation is in primary motor or in premotor cortex. A myeloarchitectonic approach, which has also become available for neuroimaging, enables us to describe some properties of cortex (Kuehn et al. 2017). Primary motor cortex, for example, is characterized by higher cortical myelin content and higher cortical thickness compared to adjacent premotor cortex and with somatosensory cortex located on the caudal bank of the central sulcus and postcentral gyrus (Fischl and Dale 2000; Glasser and van Essen 2011; Lutti et al. 2014). Describing the anatomical parcels underlying the larynx representations can inform evolutionary hypotheses and provide clues about their functional relevance.

This study sought to determine the anatomical location of larynx-related neural activity in individual subjects and to characterize the cortical structure underlying these representations. Our experimental design aimed to isolate brain activity related to voice production by controlling for breathing-related movements and movements of articulators. In one task, we identified the (supralaryngeal) articulation and the (laryngeal) vocalization component of speech during syllable production using a factorial design described in a previous study (Murphy et al. 1997). We refer to the latter ‘vocalization’ component as an index for laryngeal activity during voice production, while other studies have referred to it as ‘phonation’ or ‘voicing’. In a second task, we localized the separate neural representations of lip, tongue and larynx using highly controlled basic movements, while breathing movements were matched across conditions.

In order to characterize the microstructural properties underlying the larynx representation, we compared their myelin content and cortical thickness derived from structural and quantitative MRI measurements. These quantifications give an indication of the type of cortex underlying the activated regions, to inform our knowledge about the organization of the human larynx motor cortex.

## 2. Materials and Methods

### Subjects

20 subjects (12 females, 18 – 40 years (27.4 ± 5.6, mean ± SD), 5 self-reported left-handers) took part in the study. All subjects were self-reported native English speakers; two were raised bilingually and three were fluent in a second language. All reported normal hearing, normal or corrected-to-normal vision, no neurological impairments and no history or diagnoses of speech disorders. The study was approved by the Central University Research Ethics Committee (CUREC, R55787/RE001) in accordance with the regulatory standards of the Code of Ethics of the World Medical Association (Declaration of Helsinki). All subjects gave informed consent to their participation and were monetarily compensated for their participation.

### Experimental design and task

We used two tasks to map the motor representations of the articulators and the larynx. Task 1 engaged the speech motor system in an ecologically valid way that required the participants to utter a short syllable sequence in different conditions. Task 2 required the participants to perform basic movements that are commonly used in other localizer studies.

Subjects practiced all tasks outside the scanner to be sure they understood the task requirements. Articulator movements and vocalizations as well as breathing were demonstrated by the experimenter and practiced until the subjects performed them as required.

#### Task 1 - syllable production task

Subjects were instructed to produce the utterance “/la leɪ li la leɪ li/” in four different conditions: speaking (overt speech), supralaryngeal articulation only (silent mouthing), overt vowel sound production of vowel /i/ six times but no articulation (vowel production), and thinking (covert speech). The vowel /i/ was chosen because it is a natural and familiar sound that requires laryngeal movement during vocalization, but involves minimal movement of the jaw muscles, lips and pharyngeal part of the tongue (Grabski et al. 2012). Subjects were instructed not to whisper during the ‘silent mouthing’ or ‘covert speech’ conditions as this would involve laryngeal motor activity.

For all conditions, the breathing pattern was explicitly instructed using the fixation symbol on the screen (Figure 1A). Subjects were instructed to inhale for 1.5 s (fixation was a square) and exhale for 4 s (fixation was a cross); the utterance in each condition was performed once during each 4-s exhalation. Each condition was performed in blocks lasting 22 s, which corresponded to four repetitions of the breathing cycle. Each block was followed by a rest period of 8 s with normal breathing (i.e. not explicitly instructed). This rest period allowed the subjects to relax their breathing patterns and to maintain a comfortable respiratory state. The four conditions were presented in a fixed pseudo-random order following a balanced Latin-square design wherein each condition was repeated five times; each condition followed and preceded each of the other conditions once and the same condition was not presented consecutively. Subjects were instructed to keep their mouth slightly open during the whole session and to keep the jaw relaxed.

**Figure 1.**
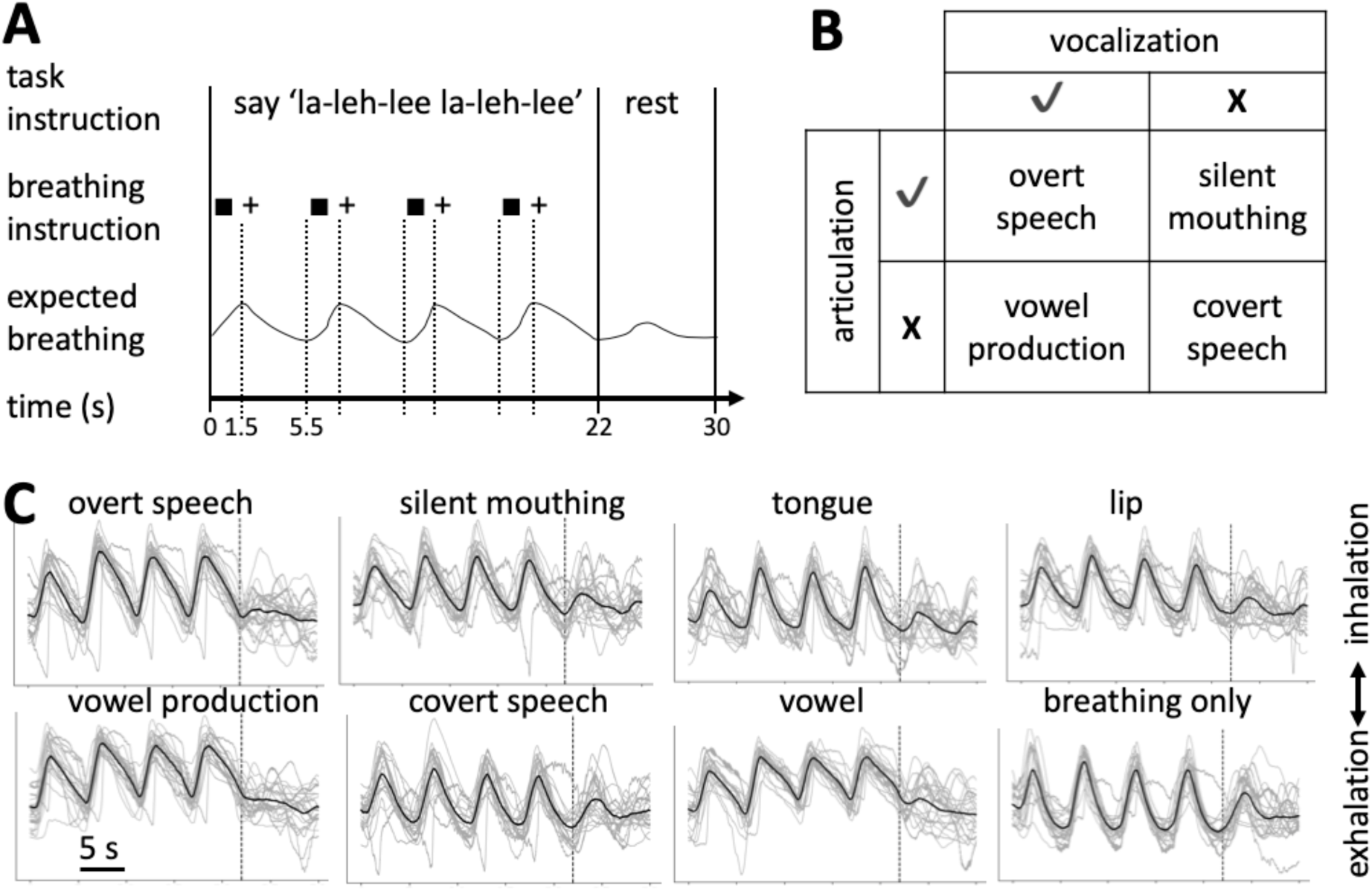
Task paradigm and breathing patterns. **A:** Timing of the task paradigm for an example block of the ‘overt speech’ condition from the syllable production task (task 1). The utterance was repeated four times while breathing was instructed followed by a resting period with normal breathing. Square – inhale, + - exhale. **B:** Factorial design used in the syllable production task. The four task conditions can be described according to orthogonal main factors for laryngeal activity during vocalization and supralaryngeal articulation. Breathing was instructed during all conditions as shown in A. **C:** Breathing traces during syllable production task (left two panels) and basic localizer task (right two panels). Black line: group mean after normalizing all individual breathing traces to the same amplitude; grey lines: average traces for individual subjects (*n* = 20). The dashed vertical line indicates the end of the 22-s task block and the beginning of the 8-s resting period with normal breathing.

#### Task 2 - basic functional localizer

There were four task conditions: tongue retraction, lip protrusion, production of the vowel sound /i/, and a ‘breathing only’ condition. The three task conditions for basic speech movements were contrasted with the ‘breathing only’ condition in which breathing was explicitly instructed as above. Articulator movements and vowel production were repeated during 4 s of exhalation at a rate of approximately 1 - 2 repetitions/s as described for task 1. Subjects were instructed to keep the movement rate constant throughout the scan for all movement types. Breathing instructions, task timing and randomization of the blocks were the same as described for task 1 above, except that each condition was repeated four times during the scan run. Note that each 22-s block was followed by an 8-s period of rest with normal breathing, as above.

### Hand localizer

During the scanning session, subjects performed the basic localizer first and then the syllable production task. In between these two tasks involving vocalizations, the subjects performed a phonological and semantic judgement task (Devlin et al. 2003). Participants had to indicate a yes/no response by pressing a button with the right index or the middle finger every 3 s. The task data was used here only to localize the hand representation in the left hemisphere; the language task data are not reported.

### MRI Data Acquisition

MRI data were obtained at the Oxford Centre for Human Brain Activity (OHBA) using a 3-T Siemens Prisma scanner with a 32-channel head coil. Two structural images of the whole brain were acquired; a T1w image (MPRAGE sequence; 1 mm^3^ isotropic resolution, TR = 1900 ms, TE = 3.97 ms, TI = 905 ms, 8° flip angle, bandwidth = 200 Hz/pixel, echo spacing = 9.2 ms, FOV = 192 × 192 × 174 mm^3^) and a T2w image (SPACE turbo-spin-echo sequence; 1 mm^3^ isotropic resolution, TR = 3200 ms, central TE = 451 ms, variable flip angle, bandwidth = 751 Hz/pixel, echo spacing = 3.34 ms, echo train duration = 919 ms, Turbo Factor = 282, FOV = 256 × 256 × 176 mm^3^, GRAPPA acceleration factor 2).

For task fMRI, whole-head T2*-weighted echo-planar images (TE = 30 ms) were acquired using multiband sequence (factor 6, TR = 0.8) at 2.4 mm^3^ isotropic resolution. Task fMRI parameters were adopted from the ABCD study (https://biobank.ctsu.ox.ac.uk/crystal/docs/brain_mri.pdf, Casey et al., 2018). Two task fMRI scans were conducted lasting 8 min (600 volumes, task 2) and 10 min (750 volumes, task 1). In between the two tasks, subjects performed a phonological and semantic judgement task for 9 min, which did not involve vocalization.

Furthermore, a multiparameter mapping (MPM) protocol was acquired (Weiskopf et al. 2013; Lutti et al. 2014). Proton density-weighted (MPM_PDw_), magnetization transfer-weighted (MPM_MTw_) and T1-weighted (MPM_T1w_) images were acquired using a tailored pulse sequence (1 mm^3^ isotropic resolution, FOV = 256 × 224 × 176 mm^3^, TR = 25 ms, bandwidth = 488 Hz/pixels, first TE/echo spacing = 2.3/2.3 ms, 6° flip angle (MPM_PDw_, MPM_MTw_) or 21° (MPM_T1w_), slab rotation = 30°, and number of echoes = 8/6/8 (MPM_PDw_/MPM_MTw_/MPM_T1w_ respectively), GRAPPA acceleration factor 2 × 2, 40 reference lines in each phase encoded direction.

Quantitative R1 (= 1 / T1) maps were estimated from the MPM_PDw_ and MPM_T1w_ images as demonstrated in Weiskopf et al. (2013), which was extended by including a correction for radio frequency transmit field inhomogeneities (Lutti et al. 2010) and imperfect spoiling (Preibisch and Deichmann 2009). Regression of the log-signal from the signal decay over echoes across all three MPM contrasts was used to calculate a map of R2* (= 1 / T2*) (Weiskopf et al. 2014). The transmit field map was calculated using a 3D echo-planar imaging (EPI) spin-echo (SE)/stimulated echo (STE) method (Lutti et al., 2012, 2010; FOV = 256 × 192 × 192 mm^3^, matrix = 64 × 64 × 48 mm^3^, TE = 39.06, mixing time = 33.8 ms, TR = 500 ms, nominal α varying from 115° to 65° in steps of 5°, acquisition time 4 min 24 s) and was corrected for off-resonance effects using a standard B0 field map (double gradient echo FLASH, 3 × 3 × 2 mm^3^ resolution, whole-brain coverage). The MPM parameter maps took approximately 20 minutes to acquire. In addition to the MRI scans mentioned here, participants underwent a diffusion-weighted scan and a resting-state scan (data not reported here). The total duration of the entire scanning session was approximately 1.5 h, preceded by approximately 45 min of briefing and task practice.

Physiological recordings were carried out throughout scanning using a respiratory belt to measure the breathing pattern and a pulse oximeter to monitor the heart rhythm during the scan. Physiological data were recorded using a Biopac MP150 (Biopac, Goleta, CA, USA) at a sampling frequency of 500 Hz. Subjects wore ear-plugs and MRI-compatible head phones (OptoActive-II, Optoacoustics Ltd, Moshav Mazor, Israel), which reduced scanner noise using electrodynamic noise-cancelling. At the beginning of the scanning session, the headphones were calibrated and noise-cancelling performance was further monitored throughout the session. Prior to each functional scan the attenuation algorithm ‘learned’ the scanner noise for 16 s. During the two tasks involving vocalizations, subjects were audio-recorded using an MRI-compatible microphone with noise cancelling (Dual Channel FOMRI-III, Optoacoustics Ltd, Moshav Mazor, Israel) at a sampling rate of 22,050 Hz.

### Behavioral data analysis

The subjects’ vocal behavior for the tasks was manually assessed using the audio recordings. The breathing patterns during the task blocks recorded using the Biopac were inspected visually to verify that subjects complied with the breathing instruction. Individual breathing traces were cropped into segments of 30 s, which consist of 22 s of instructed breathing during the task block and 8 s of subsequent rest period with normal breathing (Figure 1C).

### Structural MRI analysis

T1w and T2w scans were pre-processed using the HCP-pipeline (Glasser et al. 2013) cloned from the ‘OxfordStructural’ - fork (https://github.com/lennartverhagen/Pipelines). The processing pipeline includes anatomical surface reconstruction using FreeSurfer and automatic assignment of neuroanatomical labels (Fischl 2012; Jenkinson et al. 2012). The T2w image was registered to the T1w image using FSL’s FLIRT using spline interpolation (Jenkinson et al. 2002).

The image of the T1w scan was divided by the image of the T2w scan to create a T1w/T2w-ratio image. The T1w/T2w-ratio was mapped onto the native midthickness surface and then resampled to the 164k standard (fs_LR) surface mesh using adaptive barycentric interpolation (approx. 164.000 vertices per hemisphere) using Workbench Command (www.humanconnectome.org/software/connectome-workbench.html). Mapping was performed with the ‘-volume-to-surface-mapping’ command using the ‘-myelin-style’ option.

This T1w/T2w map has been shown empirically to correlate with cortical myelin content (Glasser and van Essen 2011; Glasser et al. 2014). In addition to T1w/T2w myelin maps, the HCP-pipeline provides automatic generation of cortical thickness surface maps.

MPM parameter maps were reconstructed and pre-processed using the hMRI-toolbox (Tabelow et al. 2019) embedded in the Statistical Parametric Mapping framework (SPM12, Wellcome Trust Centre for Neuroimaging, London, UK). For one subject, MPM data were excluded due to motion-induced blurring. MPM_MT_, MPM_R1_ and MPM_R2*_ maps were registered to the MPRAGE T1w scan using FLIRT rigid-body transformation and spline interpolation and then mapped to the surface using the same steps as for the T1w/T2w myelin map. The three MPM parameter maps have been shown to correlate to different degrees with myelin content in white matter, subcortical structures and grey matter (Draganski et al. 2011; Callaghan et al. 2014; Lutti et al. 2014; Bagnato et al. 2018). Finally, one step of surface-based smoothing (FWHM = 4 mm) was applied to the five surface maps of interest - T1w/T2w map, three MPM parameter maps and cortical thickness map.

### fMRI data analysis

Functional MRI data processing was carried out using FEAT (FMRI Expert Analysis Tool) Version 6.00, part of FSL (FMRIB’s Software Library, www.fmrib.ox.ac.uk/fsl). The following pre-statistics processing was applied: motion correction using MCFLIRT (Jenkinson et al. 2002); non-brain tissue removal using BET (Smith 2002); spatial smoothing using a Gaussian kernel (FWHM = 5 mm); grand-mean intensity normalization of the entire 4D dataset by a single multiplicative factor; low-frequency drifts were removed using a temporal high-pass filter with a cut-off of 90 s for all three tasks. Motion corrected images were unwarped using a fieldmap and PRELUDE and FUGUE software running in FSL (Jenkinson 2003). Registration to the high resolution structural scan and standard 2 mm MNI-152 template was carried out using FLIRT (Jenkinson and Smith 2001; Jenkinson et al. 2002). Registration from high resolution structural to standard space was then further refined using FNIRT nonlinear registration (Andersson et al. 2007).

Time-series statistical analysis was based on a general linear model (GLM) implemented in FILM with local autocorrelation correction (Woolrich et al. 2001). Standard motion correction parameters and individual volumes that were motion outliers, determined using fsl_motion_outliers, were included as separate regressors at the first level for each subject.

For the syllable production task (task 1), the conditions were analyzed in a factorial model that allowed the (supralaryngeal) articulation and the (laryngeal) vocalization component of the task to be separated (Figure 1B). Brain activity associated with the control of articulation was defined as (‘overt speech’ *minus* ‘vowel production’) *plus* (‘silent mouthing’ *minus* ‘covert speech’) and the main contrast for vocalization was derived by the contrast (‘overt speech’ *minus* ‘silent mouthing’) *plus* (‘vowel production’ *minus* ‘covert speech’). The 8-s periods of rest with normal breathing between condition blocks served as baseline and were not modelled as a separate explanatory variable in the GLM; they were not included in contrast to any task condition in the analysis of this factorial design.

For the basic localizer task (task 2), activity during each condition for basic speech movements was assessed relative to the ‘breathing only’ condition with instructed breathing. The periods of rest with normal breathing in between conditions served as baseline (i.e. they were not modelled in the GLM as described for task 1). For the hand localizer task, we derived a contrast of all task conditions involving button presses relative to the rest blocks (note, the breathing instruction was not used in the hand localizer task).

All contrasts reported in the results for both tasks were assessed relative to conditions with matched breathing. In addition, we report results of all individual task conditions relative to the resting condition with normal breathing in a supplementary figure (Figure S1).

### Volumetric group average activation maps

To obtain volumetric group average maps, each individual’s statistical maps were transformed to standard space (MNI152) using a nonlinear registration. Volumetric group mean activation maps were obtained using mixed effects in FLAME (FMRIB’s Local Analysis of Mixed Effects, (Woolrich et al. 2004, Stage 1 only), where subjects are treated as random effects. Group-level *z*-statistic images were thresholded using a voxel-wise threshold of *z* > 3.5 (*p* < 0.00025, uncorrected). Note that the corrected threshold for a whole-brain analysis of these data is *z* > 5. Rather than use a small volume correction and ROI masking, we chose to display results at the lower threshold to better visualize the activity in the whole brain. The figures of the volumetric group-level results (Figure 2, 3) are focused, i.e. centered, on the voxel of maximal *z*-value within the right hemisphere central somatomotor strip. For the vocalization contrast (Figure 2) and the vowel production condition (Figure 3), we generated two separate figures focused on the right dorsal and the ventral larynx representation.

**Figure 2.**
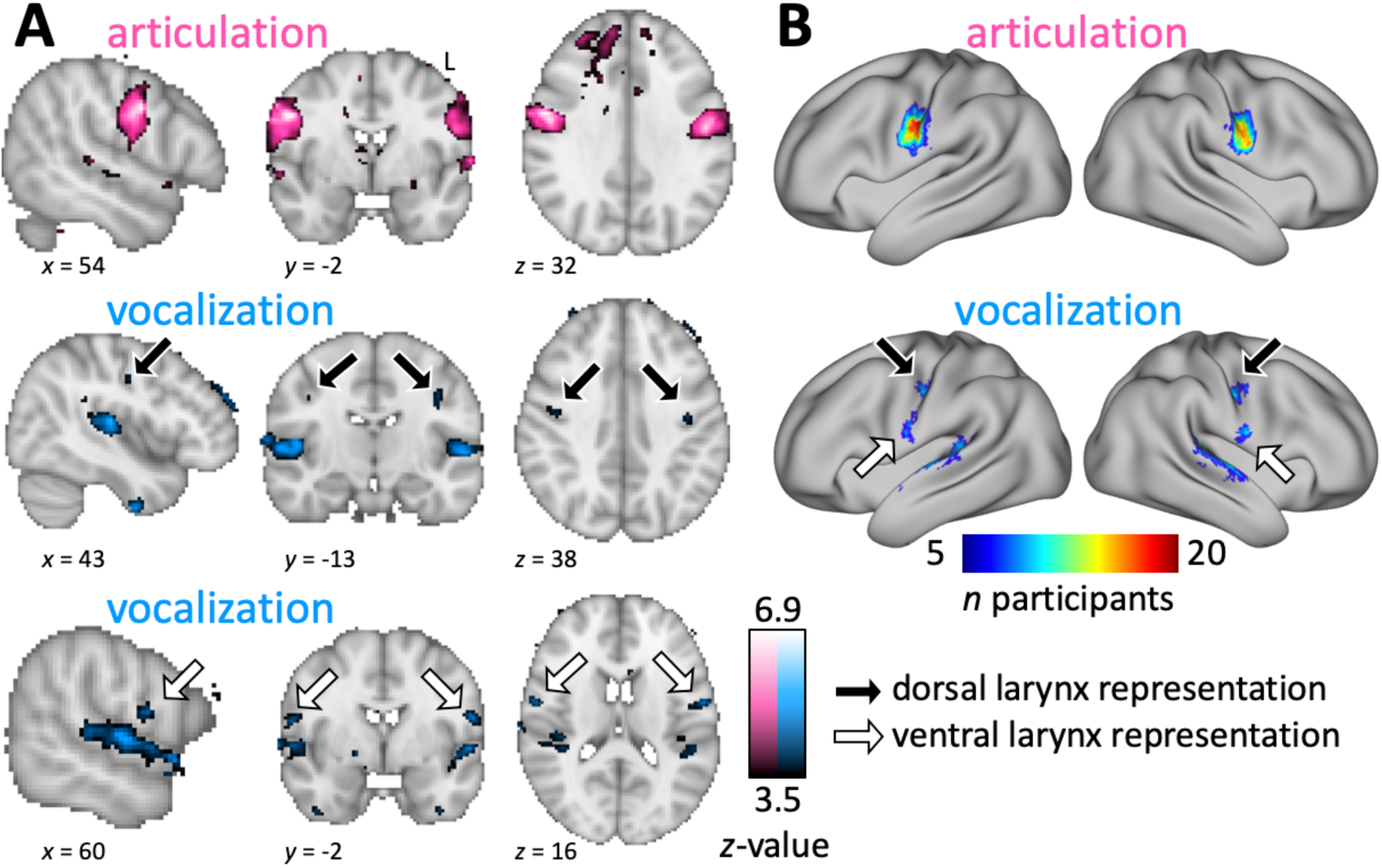
Syllable production task. **A:** Whole-brain group activation maps showing areas activated during articulation and vocalization (voxel-wise threshold *z* > 3.5, *n* = 20). For the main contrast for vocalization (blue), both a dorsal and a ventral representation are shown in separate panels. Black arrow: dorsal larynx representation; white arrow: ventral larynx representation. **B:** Surface group count maps of the same contrasts. Individual surface maps were thresholded at *p* < 0.05 (corrected voxel-wise), binarized and resampled onto the fs_LR surface. These were summed across the group of 20 subjects and thresholded at *n* > 4. Note: in both analyses, vocalization also resulted in activation of the superior temporal gyrus, presumably reflecting auditory stimulation due to hearing oneself vocalizing.

**Figure 3.**
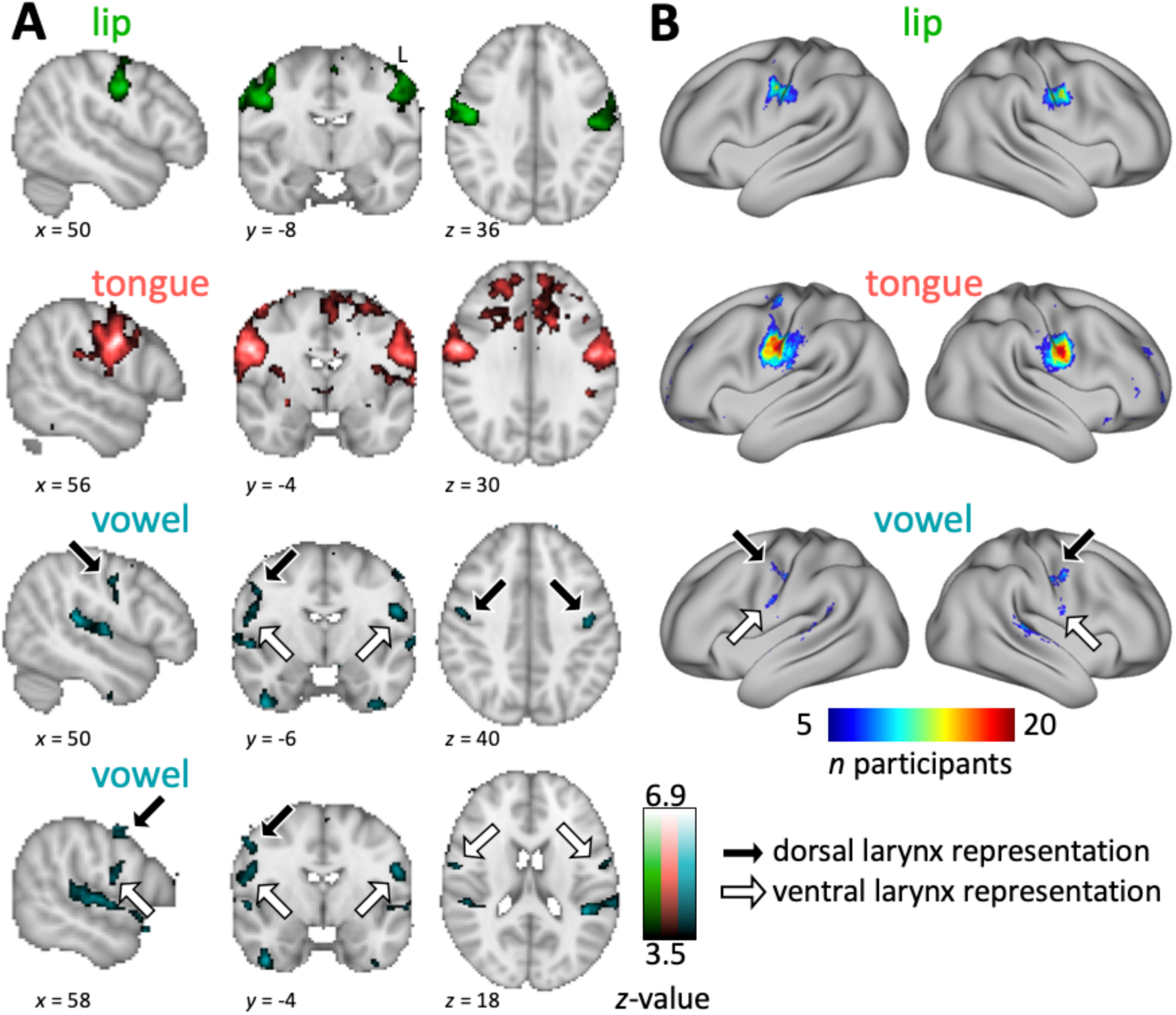
Basic localizer task. **A:** Whole-brain group activation maps showing areas activated during lip and tongue movement and during vowel production (vowel-wise threshold *z* > 3.5, *n* = 20). For the vowel production condition (turquoise), both a dorsal and a ventral representation are shown in separate panels. See legend to Figure 2 for details. **B:** Surface group count maps of the same conditions. See legend to Figure 2 for details.

### Surface group count maps

In addition to volume-based analysis, task MRI activations were assessed using surface-based analyses. Surface space permits a better visualization of cortical brain activations. Furthermore, this allowed us to perform surface-based quantifications of cortical microstructures.

Each subject’s *z*-statistic images were thresholded voxel-wise at *p* < 0.05 (corrected using a *z*-value determined based on data smoothness and the RESEL count). This thresholded map was projected onto the individual’s native midthickness surface using the ‘-ribbon-enclosed’ option in wb_command ‘-volume-to-surface-mapping’. To derive group-level surface count maps, all individual subject thresholded maps were resampled to the 32k standard (fs_LR) mesh based on the FreeSurfer registration, binarized and then summed at each vertex (Barch et al. 2013). The group count maps are shown on an inflated average brain surface, thresholded at *n* > 4 subjects. They give an indication of the between-subject variability in the areas activated by task, providing complementary information to maps of activity averaged across participants.

### Individual surface activation maxima

Surface-based activations were studied further using an ROI-based approach to isolate and focus on activations in the central somatomotor strip. The two tasks provided complementary results for further analysis with Task 1 serving as a robust localizer for the larynx representations, and Task 2 providing an accurate localizer for lip and tongue representations. In order to assess intra-individual spatial variability of the motor representations, we derived the location of individual activation maxima in both hemispheres from selected task contrasts: For the larynx, we used the main contrast for vocalization during the syllable production task (task 1). For lip and tongue, we used the task contrasts from the basic localizer task (task 2) and for the hand, we used the hand localizer task (left hemisphere only).

Different ROI masks were used for the different motor representations based on individual anatomy. In short, we used an ROI of the whole central sulcus for hand, lip and tongue, a more limited portion of the central sulcus ROI for the dorsal larynx representation and a manually defined ROI for the ventral larynx. ROI definitions are described in more detail in the supplementary material.

Individual volumetric ROIs were linearly transformed from FreeSurfer’s anatomical to functional space of the respective task fMRI scan. Within the ROI, the voxel of maximal intensity was determined from the uncorrected *z*-statistics image. It should be noted that for some subjects this local maximum did not achieve the corrected voxel-wise significance threshold, which was used to create the surface count maps (left hemisphere: hand *n* = 3, dorsal larynx *n* = 6, ventral larynx *n* = 5; right hemisphere: dorsal larynx *n* = 5, ventral larynx *n* = 4). Using a lower uncorrected threshold is justified given our goal to visualize and assess spatial variability of the activation maxima. Activation maxima were manually inspected in the subject’s native volume space to confirm that the systematic approach described below captured task-related activations. The activation maxima were mapped to the individual’s native midthickness surface, resampled to the 32k standard (fs_LR) surface mesh using the FreeSurfer registration, smoothed (FWHM = 1 mm), and binarized to form a small circular patch.

Given that the two tasks had some conditions in common, we were able to examine within-subject reliability of the location of activation maxima for larynx and tongue as described above compared with those derived based on the vowel condition and the main contrast for articulation. We computed the geodesic distance on the individual’s native midthickness surface for pairs of maxima (tongue condition and main contrast for articulation; main contrast for vocalization and vowel condition).

### Cortical surface features at activation peaks

We described the cortical microstructure at the individual activation peaks based on different surface measures. In order to assess cortical myelin content, we used the T1w/T2w map and the MPM parameter maps (MPM_MT_, MPM_R1_ and MPM_R2*_). We extracted the mean value for each individual subject’s surface map within the area defined by the circular patch around the vertex of maximal activation for hand, lip, tongue, dorsal larynx and ventral larynx representation, i.e. at the peaks shown in Figure 4A. These measures were obtained from the subject’s native surface mesh prior to resampling to the standard mesh. In order to account for the different ranges in intensities and to allow direct comparison of the maps, we computed *z*-scores based on the distribution of individual values in each map and each ROI. We assessed the differences in *z*-scores across the different ROIs using a linear mixed effects analyses as implemented in R’s *lmer* function (Bates and Sarkar (2007), Core Team and Foundation for Statistical Computing). The model included fixed effects for ROI (dorsal larynx, lip, tongue, ventral larynx), hemisphere (left, right) and map (T1w/T2w, MPM_MT_, MPM_R1_, MPM_R2*_) and random effects for subject. A Shapiro-Wilk test revealed a normal distribution of the data at a significance level of *p* > 0.001.

**Figure 4.**
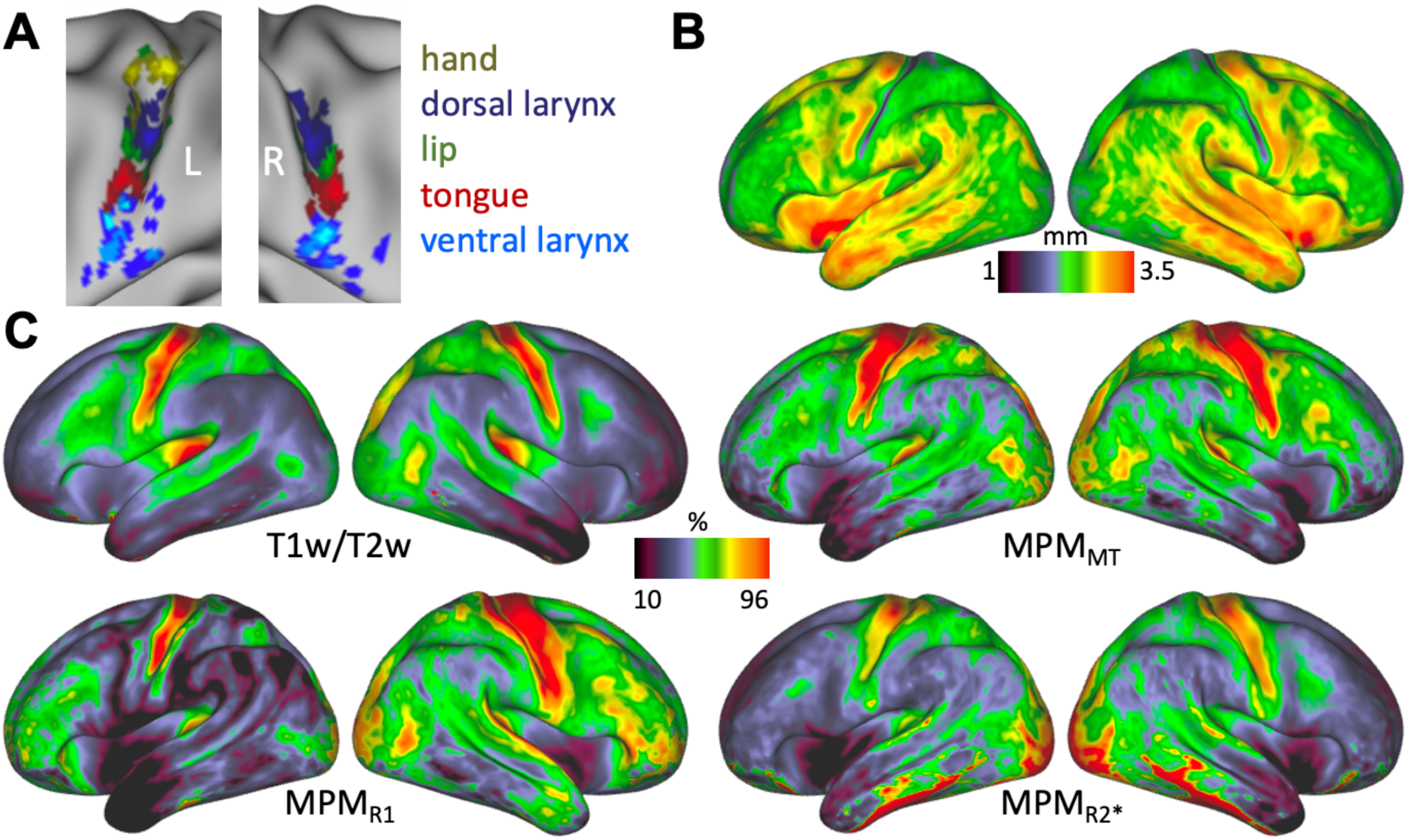
**A:** Individual activation maxima derived for hand movement (only left hemisphere), lip movement, tongue movement and vocalization (i.e. larynx activity during voice production) (*n* = 20). For vocalization, activation maxima in two separate brain regions were derived to localize the dorsal and ventral larynx representations. **B:** Average cortical thickness map (*n* = 20). **C:** Average T1w/T2w myelin map (*n* = 20) and MPM parameter maps (*n* = 19).

In addition to cortical myelin content, we also quantified cortical thickness values at the same individual surface peaks. To increase sensitivity of the quantification described above, we excluded activation maxima that were located in a part of cortex with lower cortical thickness and thus likely to be activity in somatosensory rather than motor cortex (Fischl and Dale 2000). We used a heuristic lower threshold of 2 mm to exclude sensory activation maxima and then re-ran the statistical evaluation of myelin-values described above. A linear mixed effects analysis for the fixed effects of ROI and hemisphere and random subject effects was performed to assess cortical thickness values after exclusion of the sensory activation maxima.

In order to further characterize the differences in myelin content across the ROIs, we used an additional quantification: We computed the pair-wise Manhattan distance across the ROIs based on a vector of the four raw (i.e. non-*z*-transformed) values in each subject after excluding the sensory activation maxima. The differences across ROIs were visualized in form of a dissimilarity matrix, where we averaged Manhattan distances within each ROI first across subjects and then across hemispheres.

A linear discriminant analysis (LDA) was run to explore which combination of *z*-transformed surface features (T1w/T2w, MPM_MT_, MPM_R1_, MPM_R2*_), best discriminated the ventral from the dorsal larynx representation after excluding sensory activation maxima. The parameters of the LDA were estimated using a singular value decomposition with no shrinkage.

## 3. Results

We acquired fMRI data in 20 subjects during performance of two tasks: (1) a syllable production task required subjects to produce “/la leɪ li la leɪ li/” overtly, mouthed silently, and covertly, and to produce the vowel /i/, (2) a basic localizer task required subjects to make small repetitive movements of the lips, tongue and larynx (vowel production); an additional task was used as functional localizer for movement of the right hand.

### Auditory recordings

All subjects vocalized as instructed during the conditions that involved vocalizations in both tasks. In all other conditions, subjects remained silent, as instructed.

### Breathing during vocalization tasks

We recorded the breathing traces using a breath belt during both vocalization tasks to confirm that the breathing patterns were comparable across different conditions (Figure 1C). As expected for all conditions in both tasks, four breathing cycles were visible in the first 22 s, during which breathing was instructed. Note that in one subject, where an extension of the breathing belt was used, the breathing trace differed in overall shape, but the breathing cycles were still visible. Examination of these figures shows that the shape of the trace and variability were similar across all conditions, but some differences were observed. For example, in the vowel production condition of the basic localizer task, exhalation was more gradual and less rapid than in the other four conditions. All four conditions in the syllable production task showed a more gradual exhalation pattern than in the basic localizer task.

### Syllable production task (task 1)

#### Syllable production task - Volumetric results

We localized the cortical activations for movement control of (supralaryngeal) articulation and (laryngeal) vocalization during syllable production (Figure 2A). MNI coordinates of activation maxima in the somatomotor strip are reported in the supplemental material (Table S1). The main contrasts for vocalization and articulation were defined based on a factorial design that combined the data from all four task conditions in the syllable production task (Figure 1B). The main contrast for articulation showed activity in the mid-portion of the central gyrus. Note that we also found spurious activity in prefrontal white matter, which we presumed was induced by task-correlated movement.

The main contrast for vocalization showed activity in two somatomotor regions in both hemispheres: one located within a dorsal region of central sulcus, and a second located in a more anterior-ventral region. Portions of the superior temporal gyrus were also activated during vocalization. This is presumed to reflect auditory perception of self-generated vocalizations. The dorsal and ventral activations were separate and did not appear to be connected.

Vocalization and articulation also activated parts of cerebellum and SMA in a somatotopic fashion. Group-level volumetric results for cerebellum and SMA in both functional tasks are described in the supplementary material (Figure S2). In SMA, one single representation for laryngeal activity during vocalization was observed, while in cerebellum, two distinct representations were found.

#### Syllable production task - Surface results

Group count maps of the syllable production task projected to the surface showed that several vertices in the mid-portion of the central sulcus were commonly activated during articulation in 19 out of 20 subjects (Figure 2B). The group count map for vocalization showed separate dorsal and ventral regions robustly activated across the group. In the main contrast for vocalization, there was greater variability in the exact location of activity above threshold and the areas activated in individuals were smaller than for the main contrast for articulation. This resulted in less overlap of activated vertices for the vocalization group maps. For some subjects, vocalization-induced activity did not reach significance in relevant brain regions, which also resulted in lower values in the count map.

### Basic localizer task (task 2)

#### Basic localizer task - Volumetric fMRI results

Brain activation during movement of the tongue and the lips and for laryngeal activity during vowel production was assessed by contrasting basic speech movements with a ‘breathing only’ condition, which was matched for controlled breathing. We found distinct activation peaks that followed the predicted somatotopic organization in the mid-portion of the central sulcus with the tongue representation more ventral than the lip representation in both hemispheres (Figure 3A) (Penfield and Rasmussen 1950; Grabski et al. 2012; Carey et al. 2017). The location of the activity during tongue movement overlapped with the result of the main contrast for articulation in the syllable production task; as previously, we found spurious activity in the prefrontal lobe white matter that we presume is movement related.

Vowel production induced activity bilaterally in a central dorsal region, a ventral region anterior to the central sulcus, and in superior temporal cortex. Within the dorsal region, activation is found both in the central sulcus and on the precentral gyrus. As for the syllable production task, activity in superior temporal cortex was presumed due to hearing oneself. In the group average map, and at the statistical threshold used, the dorsal and ventral larynx representations appeared to be connected, with residual activity along the ventral central sulcus.

#### Basic localizer task - Surface results

Group count maps of the basic localizer task projected to the surface revealed the same somatotopic activity pattern as seen in the volumetric results for group average activity (Figure 3B). The tongue condition showed highly consistent activation in the mid-portion of the central sulcus during articulation (maximum overlap was achieved for all 20 subjects). The result for tongue movement was highly similar to the result of the main contrast for articulation, which is in line with the volumetric results. Complete overlap of activated vertices was not achieved for the lip condition, but the activated region was still highly consistent across subjects (maximum: 15 subjects).

For the vowel production condition, the group count maps show a dorsal and a ventral cluster, similar to the pattern seen during the main contrast for vocalization (task 1). The dorsal cluster extended from the central sulcus to the precentral gyrus and in the right hemisphere a distinct dorsal gyral and a dorsal sulcal activation can be observed. We presumed that this dorsal gyral activity, but not the sulcal activity, is a residual breathing-related effect, because it overlaps with activity from the ‘breathing only’ condition (Figure S1). The ventral activation cluster appeared to extend into a part of cortex where we expected the tongue representation, and may reflect residual tongue activity during vowel production. In contrast to the volumetric results for the group averaged activity, the count maps based on individually thresholded activation maps show the dorsal and ventral representation activated by vowel production to be clearly separate.

As described above for the main contrast for vocalization, the values in the group count maps for vowel production indicate that the location of this activity is less consistent than for the other conditions; the area activated during vocalization is also smaller, both dorsally and ventrally, than for the articulators. The highest overlap was achieved in the right dorsal larynx representation in 11 subjects.

### Individual surface activation maxima

In order to characterize the variability of brain activity across subjects, we derived individual activation peaks for the hand movement (based on the hand-localizer, only in the left hemisphere), lip and tongue movement (based on the basic localizer task) and vocalization (based on the syllable production task). The main contrast for vocalization was used to localize the dorsal and a ventral larynx representation. Figure 4A shows the spatial distribution of individual activation peaks on an inflated brain surface. Overall, the location of the peaks for the different movement types is highly consistent with similar cross-subject variability. For the ventral larynx representation, particularly in the left hemisphere, however, the location of the maxima appears to be more variable.

Within-subject reliability of the activation maxima across the two tasks was compared for ‘vocalization’ and ‘vowel production’ for both dorsal and ventral larynx representations as well as for ‘articulation’ and ‘tongue movement’. Reliability was high with a median distance across subjects of less than 10 mm for the three ROIs in both hemispheres.

### Cortical surface maps of microstructural features

We derived whole-brain average surface maps for different measures related to cortical microstructure (Figure 4B, C). Overall, the T1w/T2w map and the three MPM parameter maps (MPM_MT_, MPM_R1_, MPM_R2*_) show a similar pattern of myelin content across the cortex (Figure 4C). In all four maps, the central sulcus as well as precentral and postcentral gyrus are characteristically high in myelin, which is considered to be a defining feature of primary motor and sensory cortex (Glasser and van Essen 2011). The location of the ventral boundary of the somatomotor cortex, indicated by a steep gradient of myelin values, slightly varies across the four maps, but this boundary is consistently more ventrally located in the right compared with the left hemisphere.

The four maps that correlate with cortical myelin (T1w/T2w, MPM_MT_, MPM_R1_ and MPM_R2*_) are sensitive to different biophysical properties of the myelin, but it should be noted that their sensitivity profiles are not completely independent (Callaghan et al. 2016). Therefore, some dissimilarities regarding the distribution of myelin along the cortex can be observed across the maps. MPM_MT_ and MPM_R1_ have a stronger signal in the frontal lobe compared to the T1w/T2w and the MPM_R2*_ map. In addition to myelin, the R2* signal is influenced by cortical properties such as iron content and calcium (Wu et al. 2009; Bagnato et al. 2018). For the T1w/Tw2 map, the underlying biophysical model is less well understood. The high R2* values in the ventral temporal lobe are likely a susceptibility artifact caused by signal loss at the air-tissue boundary.

In the cortical thickness map, a prominent strip of low values (i.e. thinner cortex) can be observed at the posterior bank of the central sulcus, indicating the location of primary sensory cortex (Fischl and Dale 2000) (Figure 4B). Cortical thickness values are high (i.e. thicker cortex) in the anterior bank of the central sulcus and in the precentral gyrus, which is indicating the location of primary motor cortex. In the cortical thickness map, the ventral boundary of primary sensory cortex can be determined by a sharp gradient at the level of the subcentral gyrus. The location of this boundary is comparable with the location described above in the other maps. The same hemispheric difference can be observed with the somatomotor boundary being located further ventrally in the right hemisphere.

The whole brain maps demonstrate that these cortical surface measures are informative about the cortical microstructure underlying our functional activation maxima, which we describe in the next section. The result of the reconstructed MPM parameter maps *per se* is not a primary result of the paper.

### Microstructural properties at activation maxima

Next, we determined the intensity value of the microstructural surface maps at the individual subjects’ activation maxima for the different movement types (Figure 5). Figure 5A shows z-transformed intensity values for T1w/T2w, MPM_MT_, MPM_R1_ and MPM_R2*_ maps at hand, lip, tongue as well as dorsal and ventral larynx maxima, i.e. at the peaks presented in Figure 4A. Mixed effects analyses with the factors ROI (excluding the hand ROI), hemisphere and map demonstrated that there were highly significant effects of ROI (*F*(3, 600) = 60.69, *p* < 0.001) and hemisphere (*F*(1, 600) = 43.54, *p* < 0.001) but no significant effect of map (*F*(3, 600) < 1, *n.s.*). The significant effect of ROI reflects that values for the ventral larynx are lower than in the other ROIs (post-hoc pairwise *t*-tests with Tukey adjustment, *p* < 0.001), but the values for dorsal larynx representation, lip and tongue were not significantly different. The main effect of hemisphere reflects higher values in the right hemisphere (*p* < 0.001). The interaction between ROI and hemisphere was significant (*F*(3, 600) = 3.93, *p* = 0.009) indicating that the difference between the ventral larynx representation and the other motor representations was greater on the right hemisphere than on the left.

**Figure 5.**
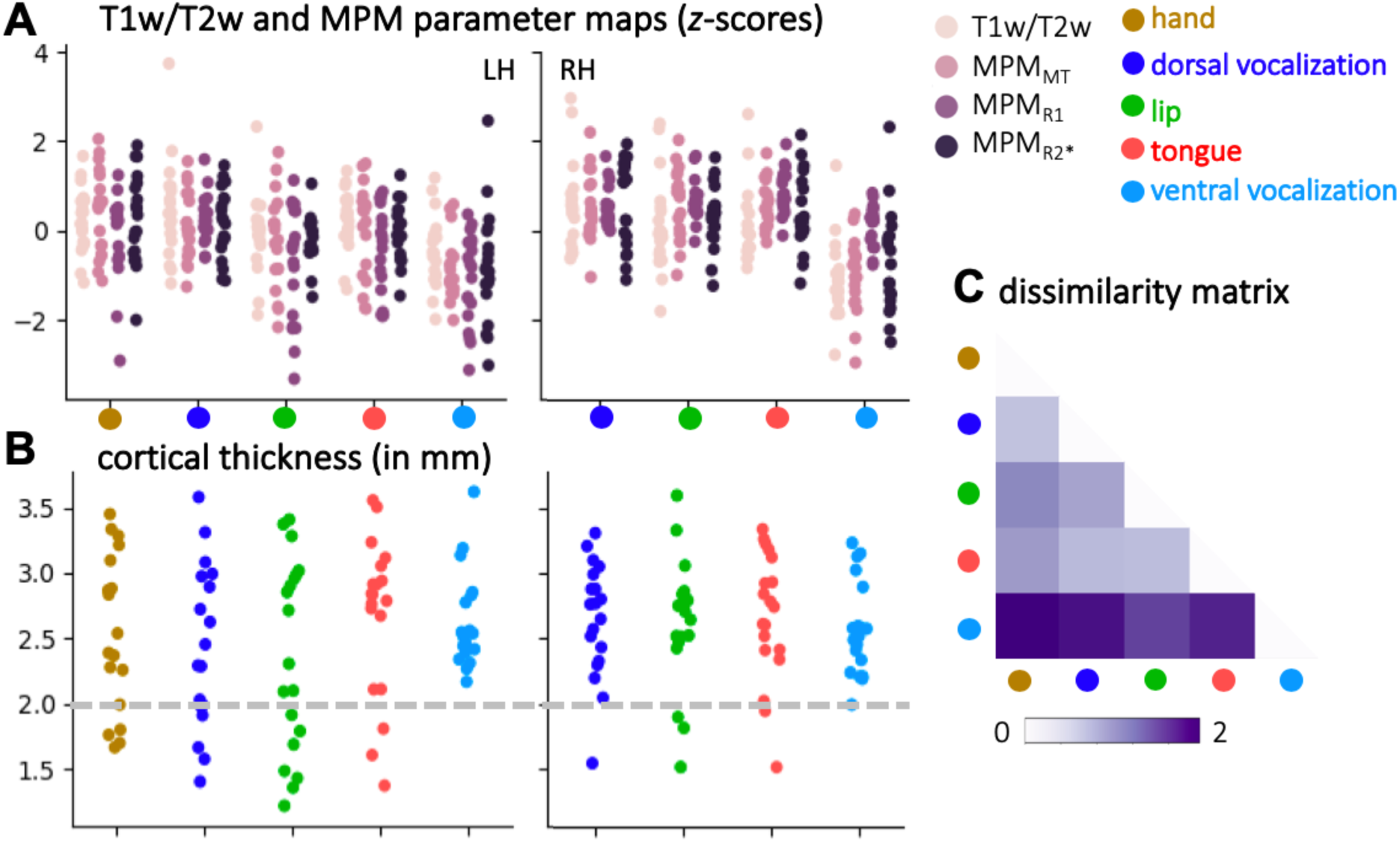
Measures of cortical microstructure derived at individual activation maxima (*n* = 19). **A:** Microstructural values that correlate with myelin (T1w/T2w, MPM_MT_, MPM_R1_, MPM_R2*_) derived at activation maxima (for individual maxima see Figure 4A). **B:** Individual values of cortical thickness. The dashed grey line indicates the threshold that was chosen to determine the exclusion of activation maxima that are presumed to be sensory due to their location in cortex < 2 mm thick. **C:** Dissimilarity matrix of the myelin values across ROIs. The matrix shows a quantitative comparison of the values shown in **A** after excluding sensory activation maxima. Pair-wise dissimilarity is defined as the Manhattan distance between each pair of ROIs.

Cortical thickness values show a high inter-individual variability for hand, dorsal larynx, tongue and lip ROIs (Figure 5B). This effect can be attributed to the fact that some maxima are located in the depth of the central sulcus, where values are low, while others are located on the anterior bank of the central sulcus, where values are high. Variability in the right hemisphere is lower, which is consistent with the location of the maxima predominantly on the anterior bank (Figure 4A). This result potentially indicates that some of derived maximal activation peaks represent activity evoked by somatosensation rather than by motor control. If maxima with a cortical thickness level below a cut-off value of 2 mm (Fischl and Dale 2000) are excluded, the comparison of cortical thickness values shows a marginally nonsignificant effect of ROI (*F*(3, 120) = 2.53, *p* < 0.061), but not of hemisphere (*F*(1, 120) < 1). The marginal effect of ROI is driven by the ventral larynx representation having lower cortical thickness values compared to the other ROIs (*p* < 0.068).

To increase sensitivity in the quantification of myelin values described above, we re-ran the linear mixed model while excluding the sensory activation maxima as defined by a cortical thickness below 2 mm. The significant main effects of ROI (*F*(3, 480) = 73.62, *p* < 0.001) and hemisphere (*F*(1, 480) = 20.81, *p* < 0.001) were stronger than reported above. As reported above, the main effect of ROI is due to decreased values in the ventral larynx representation and the hemispheric effect due to increased values in the right hemisphere. The interaction effect of ROI and hemisphere (*F*(3, 480) = 2.26, *p* = 0.08)) was no longer significant, indicating that the difference between ventral larynx representation and the other motor representations did not differ between hemispheres.

To assess effects based on the non-transformed values of the four maps (T1w/T2w, MPM_MT_, MPM_R1_, MPM_R2*_) rather than the *z*-scores, we also derived a ‘dissimilarity matrix’ based on the pair-wise Manhattan distances between ROIs after excluding the sensory activation maxima (Figure 5C). The values were averaged across hemispheres, because no significant interaction effect was found. The dissimilarity matrix reflects the effect of the ventral larynx representation being most dissimilar from the other motor representations based on these quantitative measures of cortical myelin.

A linear discriminant analysis (LDA) revealed that the following equation can be used to discriminate the ventral larynx representation from the dorsal larynx representation:

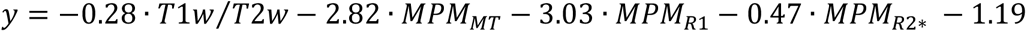

When inputting the *z*-transformed values of the surface maps at a specific location, a resulting value of y > 0 indicates that the profile of values is more similar to the ventral larynx representation than to the dorsal larynx representation. The formula indicates that MPM_MT_ and MPM_R1_ are the most informative measures to discriminate the two larynx representations (largest weighting). The mean performance accuracy of the classifier is 0.94 (while 0.5 indicates chance level performance).

Taken together, these quantifications show that the ventral larynx representation is located in cortex that has lower myelin content and lower cortical thickness compared to primary motor cortex, where the other movement representations including the dorsal larynx representation, are located.

## 4. Discussion

The goal of this study was to characterize the cortical microstructure underlying the laryngeal representations in the human brain. We localized brain activity evoked by voluntary control of laryngeal movements during vocalization using a novel paradigm. We showed that even when controlling for breathing, vocalization elicits brain activity in two separate parts of the somatomotor cortex: a dorsal region in the central sulcus and a ventral region close to the Sylvian fissure. On an individual level, the laryngeal activations and the activations during movement of hand, lips, and tongue show a consistent somatotopic arrangement. Characterization of cortical microstructure based on structural and quantitative MRI shows that the dorsal larynx representation has a similar profile to other movement representations primary motor cortex, while the ventral larynx representation has lower myelin content and cortical thickness. These results suggest that the dorsal larynx representation is the primary locus of laryngeal motor control in primary motor cortex, while the ventral larynx representation relates more to secondary activations in non-primary motor cortex.

Our experimental task was designed to separate laryngeal activity from that evoked by supralaryngeal articulation and breathing. We were able to dissociate the supralaryngeal and laryngeal components of syllable production using a factorial design and orthogonal task contrasts (Murphy et al. 1997). Covert speech is known to engage similar somatomotor brain areas to overt speech production (Kleber et al. 2007). For our purpose, however, using a ‘covert speech’ condition was essential to allow us to construct a fully factorial model aimed at isolating activity related to the execution of movements involved in articulation and vocalization while controlling for other processes such as selection and planning of movement sequences. The result of a factorial design is statistically more robust than a subtraction design, because the main contrasts include data from all task conditions improving the mathematical estimates and associated statistics. The main contrast for vocalization showed some residual activity of the tongue representation. Using the neutral vowel schwa or glottal stops instead (Gick 2002; Loucks et al. 2007; Brown et al. 2008; Grabski et al. 2012; Belyk et al. 2018), however, might have caused pharynx activity that would have been more difficult to dissociate.

In addition to the syllable production task, we used a basic localizer task that involved movement of the lips, and tongue, and vowel production. Results of the basic localizer task were consistent with the syllable production task, and within-subject variability across the comparable contrasts was low. The areas activated for the main contrast of articulation overlap with the result from the tongue localizer, which is expected given that the syllables produced mainly rely on tongue movement. The areas activated for the main contrast for vocalization overlap with the result from the vowel production condition, but some differences could be observed. Residual tongue activity resulting from producing the vowel /i/ is more apparent in the vowel production condition, suggesting that this was better controlled for in the factorial design.

When studying vocalizations, control for breathing is essential given that human vocal speech sounds are mostly produced during exhalation. Several previous neuroimaging studies, however, did not control for breathing. Some previous studies used an instructed exhalation condition for comparison with vocalization (Loucks et al. 2007; Galgano et al. 2019), but a difference in activation of regions in motor cortex was not found consistently. One likely explanation is that explicitly instructing subjects to exhale might engage laryngeal muscles in such a way that no difference to laryngeal activity during vocalization can be observed. In this study, the breathing pattern and rate were matched for all task conditions. Inhalation and exhalation were explicitly instructed on a screen and monitored using a breath belt. Although slight deviations in the shape of the breathing trace can be observed, the overall breathing traces were comparable across subjects and conditions. The deviations in breathing patterns observed in the basic localizer task suggest that breathing is less well controlled for compared with breathing during the syllable production task, where subjects produced the same utterance in different conditions.

Our results suggest that controversial findings within the neuroimaging literature can largely be explained by differences in experimental design. Several studies focused on the dorsal region as location of the ‘laryngeal motor cortex’ or ‘larynx phonation area’ (Sörös et al. 2006; Brown et al. 2008; Kleber et al. 2013; Belyk and Brown 2014; Belyk et al. 2018). Brown et al. (2008) suggested the presence of a ‘dorsolateral’ larynx representation on the crown of the precentral gyrus (*x* = 50, *y* = −2, *z* = 37) and a ‘ventromedial’ larynx representation in the central sulcus (*x* = 44, *y* = −8, *z* = 34). Our results suggest, however, that activity in the gyral portion of the dorsal region is a residual breathing-related signal and not specific for laryngeal activity during vocalization. When contrasting the ‘breathing only’ condition, during which breathing was explicitly instructed, to the resting baseline with normal breathing, we found activity in a dorsal gyral region (*x* = 55, *y* = 0, *z* = 43), which is presumably related to voluntary control of the diaphragm during instructed inhalation and exhalation (Figure S1B) (Ramsay et al. 1993; Olthoff et al. 2008). Similarly, we found the dorsal gyral activation when contrasting the ‘covert speech’ condition, which involved instructed breathing, but with no overt movement, to the resting baseline with normal breathing (Figure S1A). We also find this dorsal gyral activity during vowel production, but only when we assess it relative to the resting baseline with normal breathing (Figure S1B). When we assess vowel production relative to the ‘breathing only’ activity, only the sulcal portion of the dorsal activity, which we also found in the main contrast for vocalization, remains (*x* = 43, *y* = −13, *z* = 38). The activation that Brown et al. (2008) refer to as ‘ventromedial’ larynx area, overlaps most closely with the region that we focus in this paper as dorsal (sulcal) larynx representation.

In addition to activation of the dorsal larynx representation, multiple neuroimaging studies reported activation in a region that we refer to as ventral larynx representation (Terumitsu et al. 2006; Olthoff et al. 2008; Grabski et al. 2012). Strong evidence for the involvement of a ventral larynx representation in laryngeal motor control also comes from electrical recording from the cortical surface during vocalization and microsimulation studies (Galgano and Froud 2008; Bouchard et al. 2013; Chang et al. 2013; Breshears et al. 2015; Dichter et al. 2018). Electrocorticography demonstrated the presence of both a dorsal and a ventral larynx representation much more consistently than the fMRI literature (reviewed in Conant et al., 2014). Direct electrical stimulation of the dorsal larynx representation causes vertical larynx movement that correlates with stimulation magnitude and evokes vocalization (Dichter et al. 2018); stimulation of the ventral larynx representation causes speech arrest (Chang et al. 2017).

In addition to our novel experimental design, this study focuses on investigating individual differences in laryngeal activation patterns. The task count maps demonstrate that laryngeal activity is much less consistent across subjects compared with other movement types. In some subjects the dorsal or ventral activity did not reach the individual significance threshold. The location of the ventral larynx representation, in particular in the left hemisphere, show a much wider spatial spread across the cortex indicating large intra-individual variability. A recent anatomical study based on the dataset presented here, showed that inter-individual morphological variability in the ventral end of the central sulcus is higher than in the rest of the central sulcus (Eichert et al. 2020). It was shown that variability in sulcal morphology can explain, in part, spatial variability in functional activation peaks. As a result of the variability and the low *z*-statistical values, group level fMRI analysis in previous studies might have failed to detect one of the larynx representations or the task paradigm was not optimized to localize the larynx representations.

The existence of two separate brain regions correlating with laryngeal activity in the central sulcus raises the question as to whether both of them are involved in motor control of the larynx. Given that the larynx area in the macaque is located in a ventral premotor area (Hast et al. 1974; Jürgens 1974; Simonyan and Jürgens 2002; Coudé et al. 2011), an evolutionary ‘migration and duplication’ hypothesis has been proposed (Belyk and Brown, 2017; Jarvis, 2019; reviewed in Mars et al., 2018). The theory states that the human ventral larynx representation migrated posteriorly during evolution, possibly into a different cytoarchitectonic area. The dorsal larynx representation is thought to have evolved as human novelty in primary motor cortex.

Additional support for a duplication theory also comes from genetic profiling analyses comparing gene expressions in song-learning birds and humans (Pfenning et al. 2014; Chakraborty and Jarvis 2015). Gene expression in the avian vocal nuclei, that are involved in vocal learning, are similar to the expression profiles in both the dorsal and the ventral larynx representation of the human brain. In the context of this more general brain pathway duplication theory, it has been recently suggested that there was an additional duplication in human vocal premotor cortex, leading to the emergence of a pre-larynx motor cortex (preLMC) just anterior to the ventral larynx representation (Jarvis 2019). The proposed genetic mechanisms for the evolutionary brain pathway duplication theory in laryngeal motor control, however, remain to be verified experimentally.

An alternative interpretation of the non-human primate literature would suggest that the species differences are quantitative rather than qualitative. While traditional electrical mapping studies failed to find a focal larynx representation in primary motor cortex in non-human primates (Bailey et al. 1950; Hast et al. 1974; Simonyan and Jürgens 2002), neuronal activity correlating with laryngeal movement was observed, for example, during swallowing in macaques (Martin et al. 1997, 1999). Furthermore, tract-tracing studies showed that the premotor larynx representation in the macaque has direct connections to primary motor cortex (Künzle 1978; Simonyan and Jürgens 2002). These findings might indicate the presence of a non-human homology of the dorsal larynx representation, which is experimentally more challenging to localize due to its smaller spatial extent.

The present study is the first to characterize the microstructural properties of cortex underlying the dorsal and ventral larynx representation. The use of high-resolution quantitative neuroimaging allows us to characterize cortical architecture noninvasively (Weiskopf et al. 2013; Lutti et al. 2014). Measures from T1w/T2w maps and MPM parameter maps (MPM_MT_, MPM_R1_, MPM_R2*_) are sensitive to different microstructural properties of the cortex, but all of them have been shown to correlate with myelin to varying degrees (Glasser and van Essen 2011; Weiskopf et al. 2013; Callaghan et al. 2014, 2015; Bagnato et al. 2018). Age-related effects can confound measures of cortical microstructure (Gennatas et al. 2017; Grydeland et al. 2019), but the effects in the regions studied are expected to be minimal.

The cytoarchitectonic area underlying the larynx representations, remains to be determined histologically. Our quantification, however, suggests strongly that the dorsal larynx representation is located in primary motor cortex, which is characterized by high myelin content and thicker cortex (Fischl and Dale 2000; Glasser and van Essen 2011). The ventral larynx representation has lower myelin content and thinner cortex indicating that it is located in a different cortical territory. Based on a cytoarchitectonic map of the human brain (Brodmann 1905), we suggest that the human ventral larynx representation is located in Brodmann area (BA) 6 (premotor cortex). This interpretation is consistent with the idea that the primate brain contains multiple laryngeal representations in different cortical areas: The human ventral larynx representation is homologous to the premotor macaque larynx representation (Hast et al. 1974; Jürgens 1974; Simonyan and Jürgens 2003; Coudé et al. 2011) and the human dorsal larynx representation is homologous to a macaque larynx representation in primary motor cortex, which receives projections from the premotor larynx representation (Künzle 1978; Simonyan and Jürgens 2002).

Many individual maxima for the ventral larynx representation were located in the subcentral part of cortex. Brodmann considered this a postcentral and therefore somatosensory cortical region (BA 43), subjacent and anterior to primary somatosensory cortex (BA 3, 1 and 2) (Brodmann 1905). A magnetoencephalography study showed that air-puff stimulation of the larynx evokes activity in a ventral region (Miyaji et al. 2014). Using fMRI, BA 43 was also found to be activated by movements of the tympanic membrane associated with changes in pressure in the oropharyngeal cavity such as those that occur during gustation and swallowing and in vocalization (Job et al. 2011). These results suggest that the ventral larynx activity could be a sensory rather than a motor representation.

The question regarding the distinct functional contributions of the two larynx representations during voice productions remains unanswered. Based on our results, we formulate a hypothesis regarding the causal role of the two laryngeal representations during voice production: We propose that, as for vocalizations in non-human primates, the ventral larynx representation is involved in basic control of the vocal folds so that a vocal sound can be produced. Rapid adduction (tensioning) and abduction (relaxation) of the vocal folds at the onset and offset of a vocalization is primarily modulated by the intrinsic laryngeal muscles (Jürgens 1974; Simonyan and Jürgens 2003). Fine motor control, which is required for pitch modulations in human speech and singing, however, also relies on the vertical movement of the larynx within the trachea, which is mediated by the extrinsic muscles. We suggest that pitch control is facilitated by the dorsal larynx representation, which is located in primary motor cortex (Kleber et al. 2013; Dichter et al. 2018; Finkel et al. 2019). Activity in both intrinsic and extrinsic laryngeal muscles, however, are tightly coupled and might not be controlled by distinct brain areas (Belyk and Brown 2014). Our hypothesis is in line with an evolutionary theory suggesting that the dorsal larynx representation is unique to the human brain (Belyk and Brown 2017). Such a functional dissociation of dorsal and ventral larynx representation during vocalization, however, still needs to be tested directly using, for example, non-invasive brain stimulation.

In sum, we used neuroimaging to localize neural activity related to laryngeal motor control during vocalization, while controlling for confounding factors such as breathing and supralaryngeal articulation. We found two activated regions for laryngeal activity during vocalization, which are in anatomically distinct brain areas. Quantification of cortical microstructure suggests that the dorsal representation, but not the ventral representation, is located in primary motor cortex. It remains open, whether and how these two representations differentially contribute to laryngeal motor control.

## Acknowledgements

N.E. is a Wellcome Trust Doctoral student in Neuroscience at the University of Oxford [203730/Z/16/Z]. The project was supported by the NIHR Oxford Health Biomedical Research Centre. The Wellcome Centre for Integrative Neuroimaging is supported by core funding from the Wellcome Trust [203139/Z/16/Z]. The work of R.B.M. is supported by the Biotechnology and Biological Sciences Research Council (BBSRC) UK [BB/N019814/1] and the Netherlands Organization for Scientific Research NWO [452-13-015]. The Authors would like to thank Martina Callaghan from University College London for the MPM sequence.

## Code Availability

Upon acceptance of the manuscript, all processing code will be made available from the Wellcome Centre for Integrative Neuroimaging’s GitLab at git.fmrib.ox.ac.uk/neichert/project_larynx. FSL tools, are available from fsl.fmrib.ox.ac.uk. Connectome Workbench is available at www.humanconnectome.org/software/connectome-workbench.html.

## Supplementary Material

### Surface count maps of all individual task contrasts

**Figure S1.**
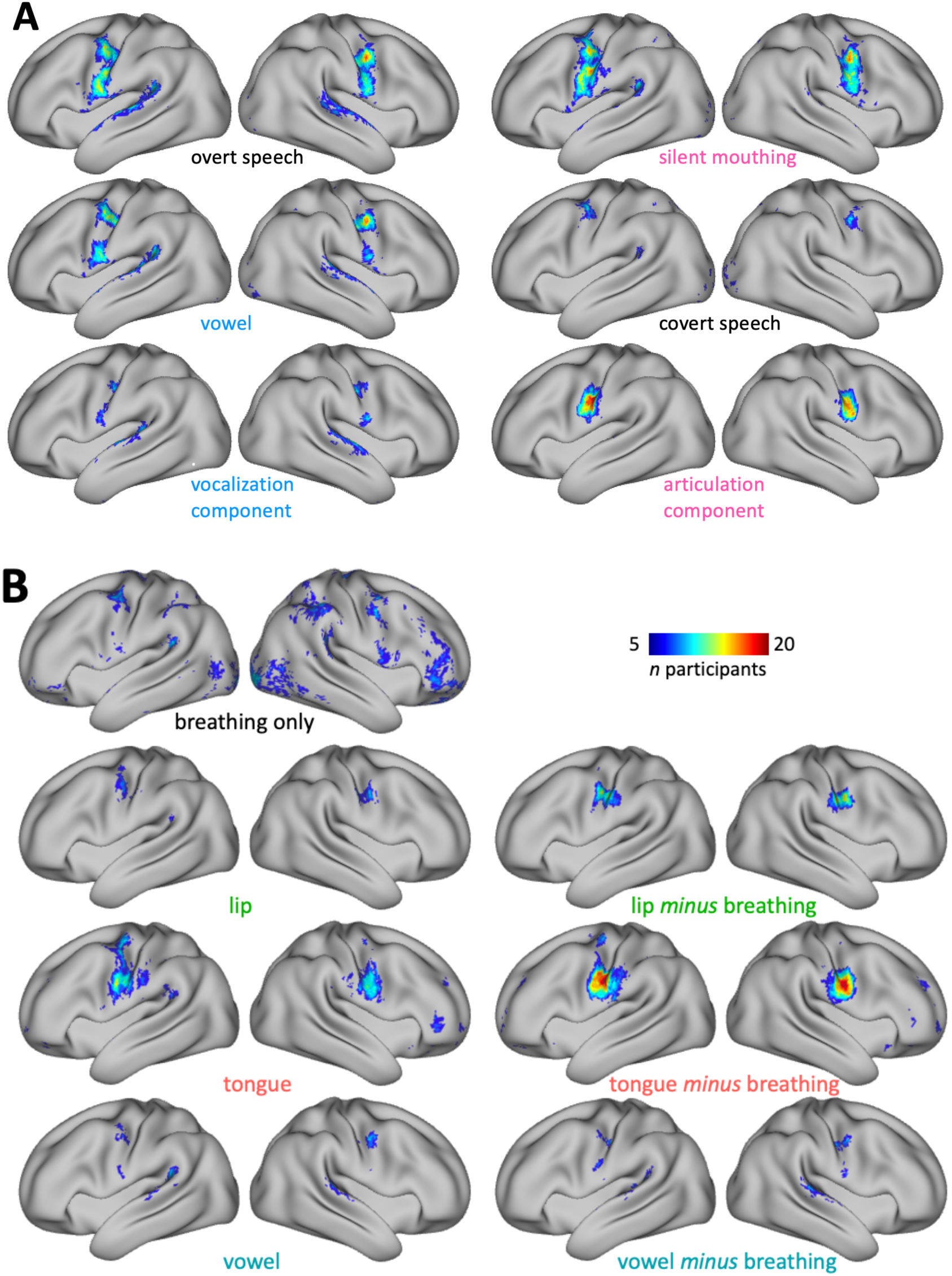
Related to Figure 2 and Figure 3. Surface count maps of all individual task contrasts. **A:** Syllable production task. Upper two rows: Four task conditions - overt speech, silent mouthing (articulation only), vowel production (vocalization only), covert speech (thinking) when compared to the resting baseline with normal breathing. Third row: Two orthogonal main contrasts for the articulation and vocalization component (these are also shown in Figure 2B). **B:** Basic localizer task. Left panel: Task conditions compared to the resting baseline with normal breathing. The ‘breathing only’ condition is a contrast between the breathing condition (where breathing was instructed) and the resting baseline with normal breathing. A lower threshold of *z* = 1.96 was used to threshold individual *z*-statistical images for this contrast due to generally lower activation. Right panel: Movement conditions compared to the breathing condition with explicit breathing instruction (these are also shown in Figure 3B).

### Task activations in supplementary motor area

The two main contrasts for articulation and vocalization - at a lower voxel-wise threshold of *z* > 2 - revealed activity bilaterally in SMA (Figure S2). A somatotopic arrangement was observed with vocalization activating a more anterior and more dorsal part of cortex. The representations of the effectors during the basic localizer task also show a clear somatotopy: In dorsal-to-ventral direction we first find the representation of the larynx, which is also most anterior, then of the lip and then of the tongue. Only one single representation of the larynx was found in SMA in both the main contrast for vocalization and in the vowel condition.

The location of the representation of the speech effectors is in line with previous accounts in the literature (Picard and Strick 1996). It has been suggested that the vertical line crossing the anterior commissure, the VAC, is a landmark for a division of SMA proper and pre-SMA (Picard and Strick 1996; Rizzolatti et al. 1996). Our results thus suggest that lip and tongue representations are located just posterior to VAC in an anterior region of SMA proper. Laryngeal activity during vocalization, however, activates a part of cortex anterior to VAC, presumably in preSMA.

### Task activations in cerebellum

Movement of the articulators and laryngeal activity also evoke activation of the cerebellum in a somatotopic fashion, which is mirroring the order observed for motor cortex (Figure S2). Most ventrally, a representation of the larynx is observed, which activates during vowel production and vocalization. This is followed dorsally by a representation of the lips and the tongue and then by a second representation of the larynx.

According to a probabilistic atlas of the human cerebellum (Diedrichsen et al. 2009), all the activations observed are located in the anterior cerebellar lobule VI. This finding is consistent with previous neuroimaging studies that found activation of the anterior-superior aspect of cerebellum during speech movements (Petersen et al. 1989; Fiez and Raichle 1997). A previous resting-state functional connectivity study demonstrated that cerebellar representations mirror the topography in cerebral motor cortex (Buckner et al. 2011). It is therefore presumed that the ventral-most representation of the larynx in cerebellum is related to the dorsal larynx representation in motor cortex and vice versa.

**Figure S2.**
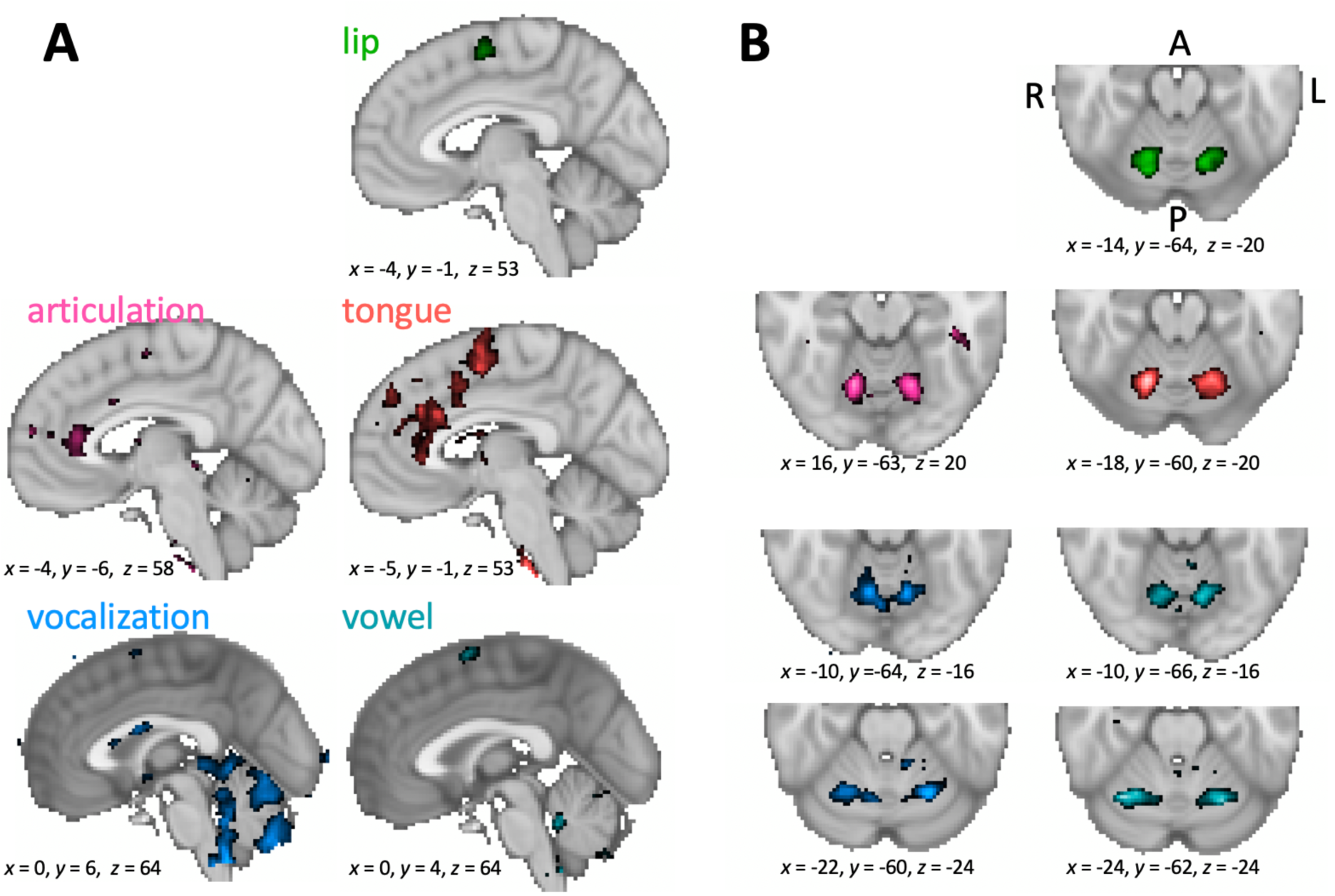
Related to Figure 2 and Figure 3. Whole-brain group activation maps showing areas activated during syllable production task (left panels) and basic localizer task (right panels) (vowel-wise threshold *z* > 3.5, *n* = 20). Coordinates are given for the voxel of maximal activation in the left hemisphere. **A:** Activity on the medial brain surface centered on the voxel of maximal activity in the left supplementary motor area (SMA) (note that for the main contrast for vocalization the threshold was lowered to *z* > 2). **B:** Cerebellar activity in the same task contrasts and shown in the same colors as in **A**. For the main contrast for vocalization and for the vowel production contrast, two separate activations are shown in different slices.

### Regions-of-interest to derive maxima during task activation

An example of the volumetric ROIs in an individual is shown in Figure S3. The central sulcus ROI used for the hand, lip and tongue was defined using FreeSurfer’s automatic volumetric labelling based on the Destrieux Atlas.

For the larynx, we identified two activation maxima in separate ROIs: One for the dorsal and one for the ventral larynx representation. The dorsal larynx ROI was a portion of the same central sulcus ROI used above from *z*-coordinates in MNI space of 50 - 30. The limits were determined empirically, so that ROI did not capture the ventral larynx representation or an unrelated supra-dorsal activation in the trunk area, which was observed in some individuals (Foerster 1931).

The ventral larynx representation lay outside the central sulcus and was located ventrally in the subcentral part of cortex. Due to the high intra-individual morphological variability in this region (Eichert et al. 2020), the ventral larynx ROI was derived manually based on individual anatomy in surface space. A liberal surface ROI was drawn on each individual’s midthickness surface covering the ventral part of the central sulcus and adjacent gyri (Figure S3A). Anteriorly, the ROI was delineated by the inferior portion of the precentral sulcus and posteriorly the ROI spanned the postcentral gyrus. If present, the lateral portion of the ascending sulcus in the subcentral gyrus was included within the ROI. The dorsal limit of the ROI was defined by a horizontal plane across the gyrus at the level of the usual location of the posterior ramus of the inferior precentral sulcus. The ventral larynx surface ROI was converted into a volumetric ROI covering the underlying cortical ribbon using wb_command. We checked that the ventral larynx ROI did not overlap with subjacent auditory cortex in the temporal lobe or inferior frontal cortex.

In some subjects, the main contrast for vocalization in the syllable production task had additional activity related to articulation of the tongue. To remove this, we transformed the coordinates for each individual’s maximal voxel from the tongue contrast (from the basic localizer task) to the functional space of the syllable production task (task 1) using rigid-body transformation and then derived a spherical ROI (7 voxels diameter) around it. This sphere was used to mask the *z*-statistic image of the main contrast for vocalization prior to localizing the maxima for laryngeal activity in the dorsal and ventral ROIs described above.

**Figure S3.**
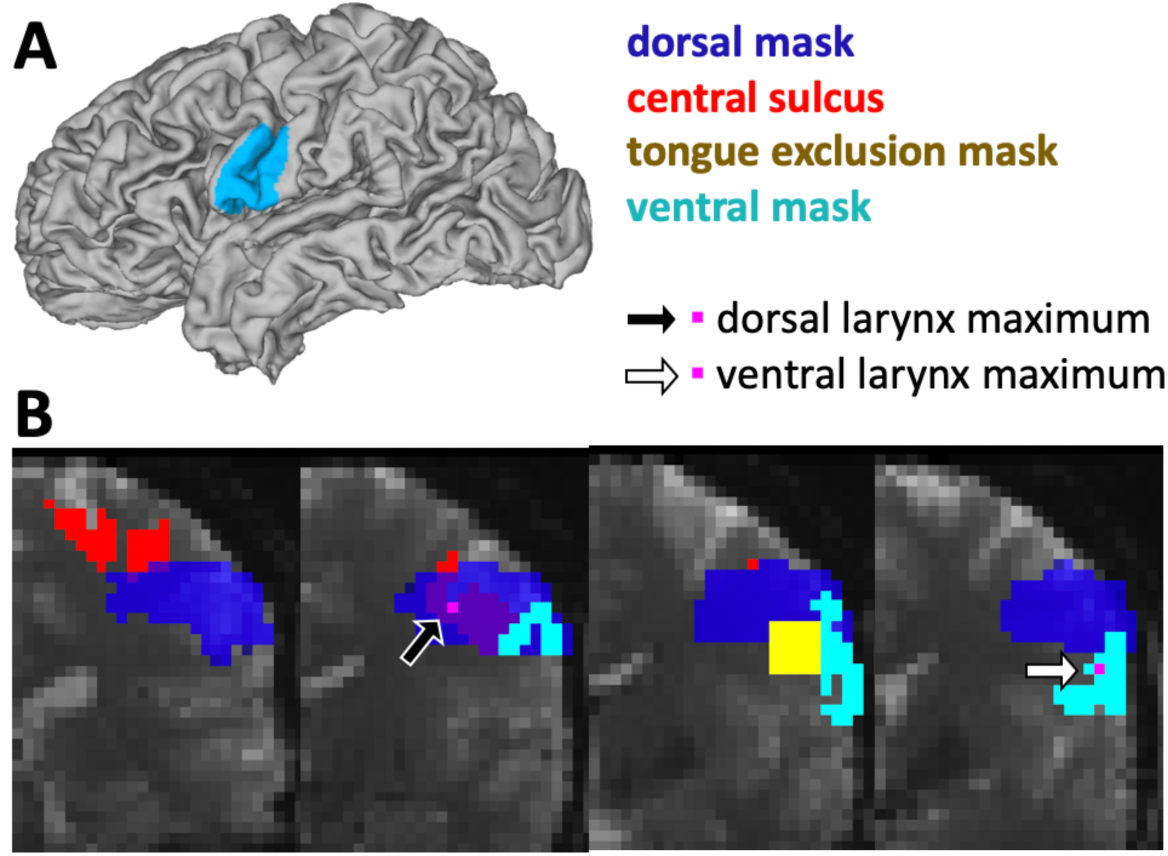
Related to Methods. **A:** Example of a manually-drawn surface ROI for the ventral larynx representation. **B:** Volumetric ROIs overlaid onto an individual’s functional scan. The central sulcus ROI (red) was used to derive the voxel of maximal activation for the hand, lip and tongue. The intersection of the central sulcus ROI and a dorsal mask (dark blue) was used to derive the voxel of maximal activation for the dorsal larynx representation (black arrow, pink voxel). The ventral mask was projected from surface to volume space (bright blue) to derive the voxel of maximal activation in the ventral larynx representation (white arrow, pink voxel). A spherical ROI around the voxel of maximal activation in the tongue contrast was masked out from the dorsal and ventral mask (yellow).

### Group-level task activation maxima

**Table S1:**
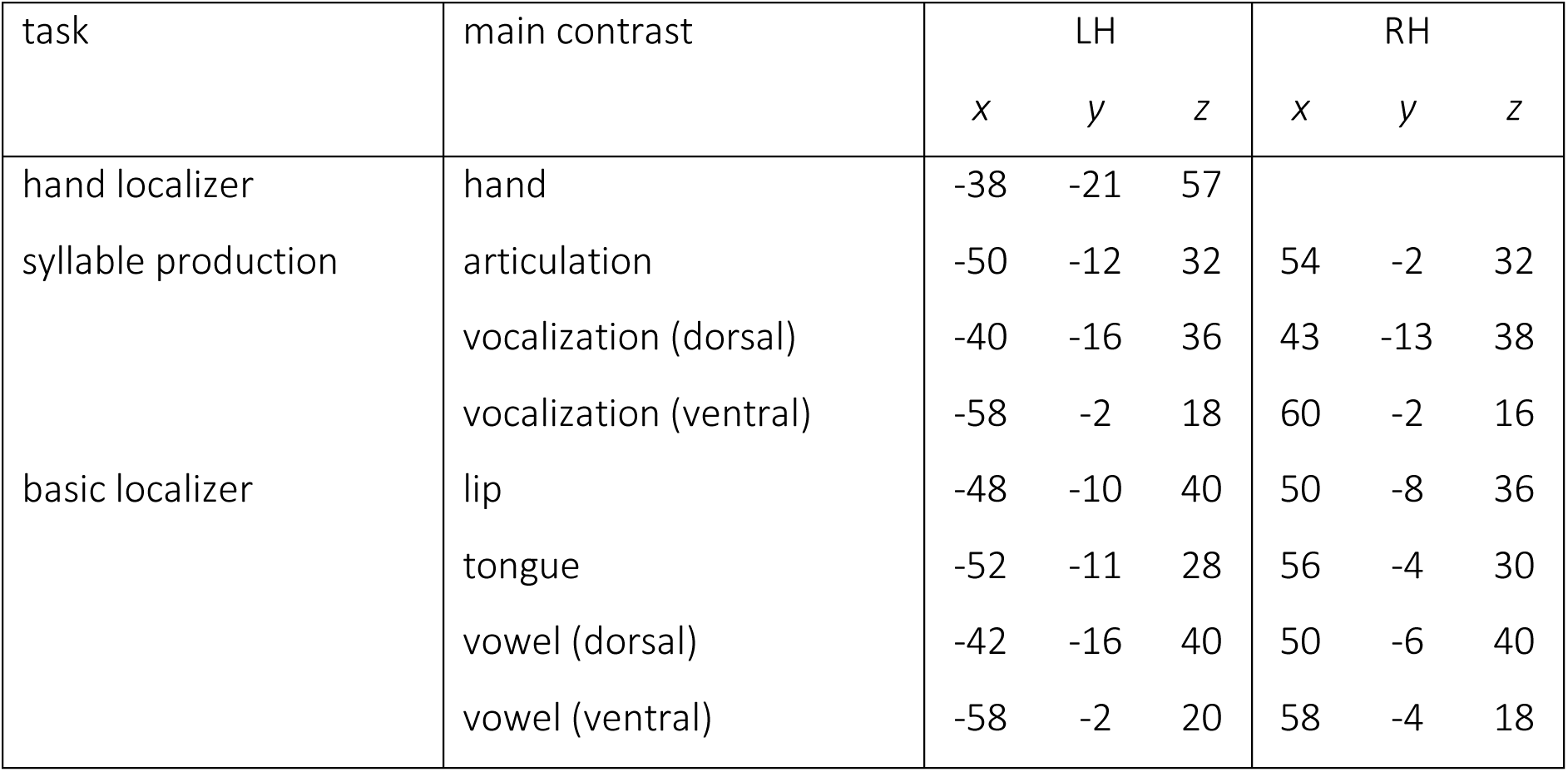
Group-level task activation maxima. Reported are MNI coordinates of maxima in somatomotor cortex for the main contrasts reported in the manuscript (LH, RH: left and right hemisphere). For the main contrast for vocalization and vowel production, a dorsal and a ventral maxima are reported separately.

